# A hand biomechanics dataset of kinematics, kinetics, electromyography, and imaging in healthy adults

**DOI:** 10.1101/2025.08.21.671503

**Authors:** Maximillian T. Diaz, Alexis R. Benoit, Kalyn M. Kearney, Troy F. Kelly, Erica M. Lindbeck, Isaly Tappan, William S. Bowers, Lavanya Durai, Justin B. Nunag, Michael B. Officer, Joel B. Harley, Jennifer A. Nichols

## Abstract

Developing musculoskeletal hand models requires a variety of experimental biomechanics data. However, collecting robust biomechanics hand data is a time intensive process leading to a lack of widely available datasets. To address this issue the biomechanics hand modeling database (BHaM) was made as a collection of experimental data to aid the development, testing, and validation of musculoskeletal models and simulations. BHaM includes two datasets: (1) a population dataset (n = 726 adults) describing hand strength (pinch and grip), self-reported hand function (Michigan Hand Questionnaire), and anthropometric measurements (from photographs), and (2) a biomechanics dataset (n = 30 adults) describing kinematics (marker-based motion capture), kinetics (isometric and isokinetic data), and electromyography (surface and fine wire) during 19 tasks across the elbow, wrist, and hand. A subset of the biomechanics dataset (n = 15 adults) also includes magnetic resonance imaging of the shoulder through wrist. Participants for both datasets were recruited to represent a diverse population of healthy adults, ranging from 18 to 91 years.

## 1 Introduction

The human hand coordinates 27 bones and over two dozen degrees of freedom any time it is in motion or exerting force.^1,2^ To achieve these feats, 30 muscles are stored in complex layers within the forearm and hand.^2^ Extrinsic muscles, like the *flexor digitorum profundus*, originate in the forearm and move the hand via multiarticular tendons, which span and create forces across multiple joints.^1–3^ In contrast, intrinsic hand muscles, like the lumbricals, are fully contained within the hand, although they can span one or more joints.^1^ This complexity makes the hand difficult to study because there are a semi-infinite number of ways to perform a single task, and the small structures make many types of *in vivo* measurements (e.g., muscle activity, joint angles, forces) difficult.^2–4^ To facilitate the study of complex structures, like the hand, musculoskeletal models (i.e., physics-based models of muscle and bones) have been developed to simulate tasks, estimate internal forces, and augment study populations.^5–8^

The dimensions of musculoskeletal models were originally developed by compiling previously published data. For example, upper limb model anthropometrics have been widely based on the United States 1988 Army Anthropometric Study, which provides population distributions for 64 unique hand and arm dimensions taken from 2,307 U.S. Army soldiers.^9^ To date, the 1988 Army Anthropometric study is still the most comprehensive public dataset on hand anthropometrics. Yet, smaller studies have shown that anthropometrics change over time and vary based on demographic characteristics, such as biological sex, age, race, and/or ethnicity. For example, <u>Jee and Yun (2016)</u> used digital calipers to measure 27 hand dimensions in Koreans, <u>García-Cáceres et al. (2012)</u> directly measured 33 hand measurements from female floriculturists, and <u>Clerke et al. (2005)</u> explored how hand shapes influenced grip strength measurements in Australian teenagers. <u>Jee and Yun (2016), García-Cáceres et al. (2012)</u>, and <u>Clerke et al. (2005)</u> all found variations in hand dimension when compared to the U.S. Army population.

Muscle properties for musculoskeletal models have primarily come from cadavers or young adults. The original models relied on published cadaveric data to determine model muscle parameters due to the infeasibility of obtaining *in vivo* measurements.^13^ While only <u>Langenderfer et al. (2004)</u> specify the age and demographics of their cadaveric specimens, it is typically assumed cadaveric specimens come from older individuals.^14–19^ Therefore, with technological improvements in medical imaging, model muscle parameters are being updated to reflect *in vivo* measurements from younger individuals.^5,8,20,21^

The development of models with parameters derived from small sample sizes is a direct result of the time required to collect and process biomechanical data and the difficulty in obtaining robust data from complex joints.^22^ Due to the complexity of obtaining hand biomechanics data, <u>McFarland et al. (2023)</u> was limited to using synthetic input data for pinch and grip tasks and comparing simulation results to previously published muscle activations. More typically, biomechanics studies recruit moderate samples (n∼30), in which the collected data is constrained to a single age demographic that may not fully represent the population. For example, <u>McFarland et al. (2019)</u> validated a musculoskeletal upper limb model using a population of 24 participants ranging from 20 to 32 years old. While this population provides strong validation for younger participants, it is unknown how the findings could change when studying older participants. Even when biomechanics studies can recruit very large samples (n > 100), the collected data does not always generalize to the population, as traditional recruitment methods in biomechanics are prone to selection bias.^23,24^

In this context, the biomechanics hand modeling database (BHaM) presented herein aims to i) provide a population dataset to understand the current hand strength, function, and anthropometrics of the population and ii) to create a biomechanics dataset of motion capture, electromyography (EMG), force recordings, and magnetic resonance (MR) images for creating musculoskeletal hand models and exploring the complex relationships between muscle activity, joint angles, and forces. For discussion regarding to what extent selection bias exists in the BHaM dataset, please see Diaz et al. (2024).

## 2 Methods

### 2.1 BHaM Population Dataset

For this IRB-approved study (UF IRB #202002061), a portable laboratory setup was designed and taken to 18 different locations across the University of Florida in Gainesville, Florida; Alachua County in North Central Florida; Brevard County in East-Central Florida; and the 2022 North American Congress on Biomechanics (NACOB) held in Ottawa, Canada. Across 41 collections, a total of 726 participants [male/female/intersex/no answer: 291/426/5/4, age range: 18 – 91 years, mean age (st. dev.): 36.6 (19.2) years] were recruited (Figure 1). The study consisted of a 15-minute testing session to record demographics, assess hand function via the Michigan Hand Questionnaire,^25^ measure grip strength using a clinical dynamometer, measure lateral pinch strength using both a clinical dynamometer and a six-degree-of-freedom time-series force sensor, and obtain a digital photograph of the palm to enable direct measurement of anthropometrics. Data were collected from each participant’s dominant hand, defined as the hand used for writing, eating, and brushing teeth.

**Figure 1.**
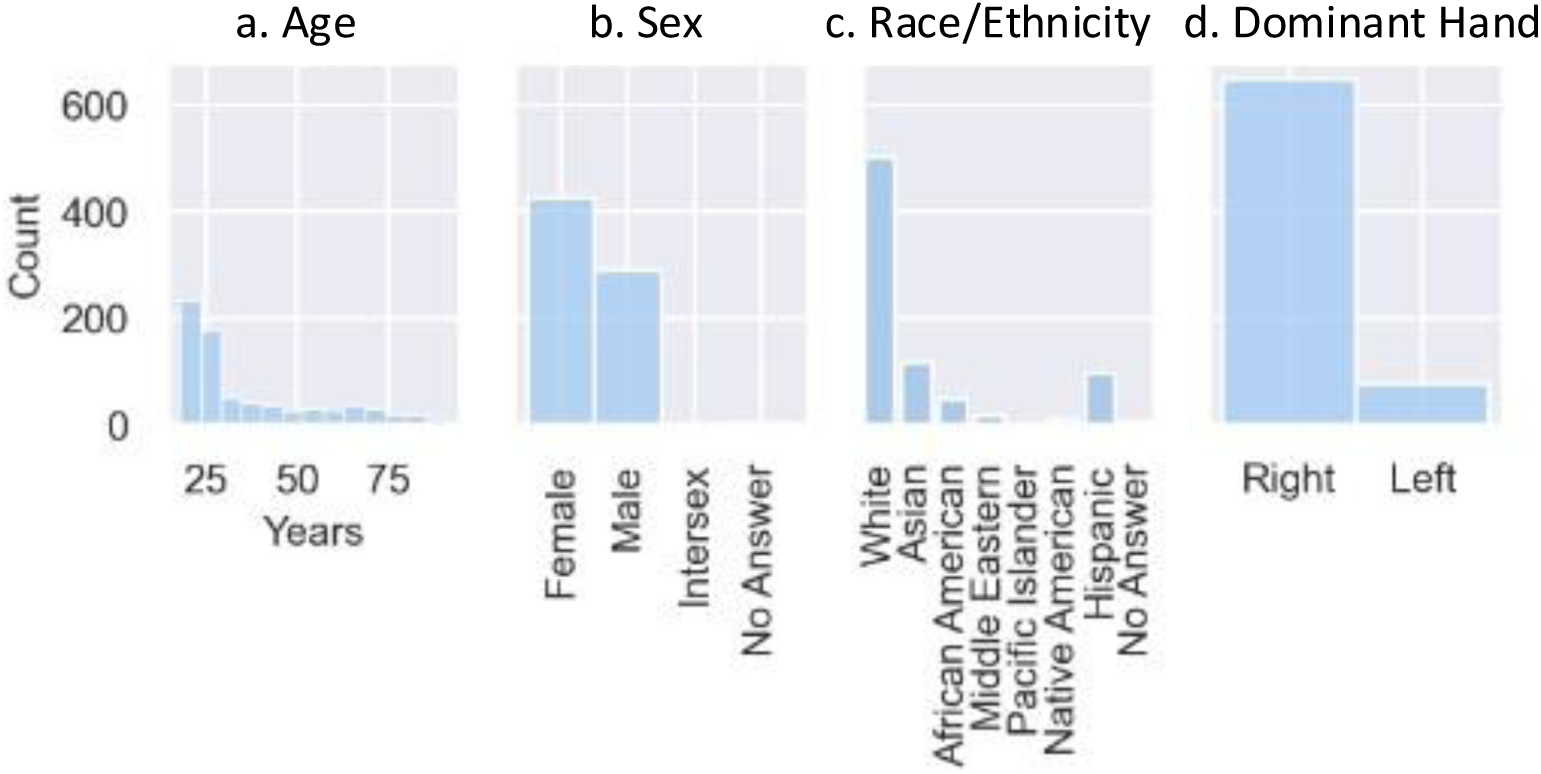
Distribution of participants within the BHaM Population Dataset. The collected population (a) is skewed towards younger individuals due to higher recruitment rates at university settings, (b) has a higher rate of females than males compared to the United States population, (c) is skewed towards white individuals, with a lower rate of Black participants than is found in the Alachua County area, and (d) a higher rate of left-handed individuals compared to the United States population.

Hand strength was assessed with both clinical dynamometry and a six-degree-of-freedom force sensor. Grip strength was recorded using a digital Jamar grip dynamometer (DHD-1, B&L Engineering, California, United States) to record a single trial maximum grip test.^26^ The dynamometer handle was set to the second position, and participants were instructed to complete the task with their arm held straight out in front of them while standing [shoulder flexion: 90°, elbow flexion: 0°, wrist supination/deviation: 0°]. This posture was chosen as it led to the most consistent positioning of participants across different locations and research team members. Lateral pinch strength was assessed with hand-held pinch gauges (10lb, 30lb, 60lb Pinch Gauges, B&L Engineering, California, United States) using a single trial maximum lateral pinch test.^26,27^ Participants were instructed to give a thumbs-up, a research team member then placed the sensor on top of the index finger, and the participant was instructed to pinch down as hard as they could. Testing was primarily done with a 30-lb rated gauge; however, if a participant failed to reach 10-lbs or exceeded 30-lbs, the test was redone with a 10-lb and 60-lb rated gauge, respectively. Lateral pinch strength was also measured using a six-degree-of-freedom, time-series force sensor (Mini40, ATI, North Carolina, United States) (100Hz); each participant completed three 5-second maximum pinch tests with 10-second rest between trials. Participants were cued to start and stop pinching with timed audio cues (“Push”, “Relax”) spoken by the computer. Visual feedback was provided to all participants in the form of the z-component of the raw force signal (downward force) on the screen. Across all strength measurements participants were verbally encouraged by research team members using standard phrasing of “Push, Push, Push”.

Hand measurements were obtained using participant hand images and photogrammetry.^9,28^ Images were taken with the participants hand supinated (palm up) and with fingers in the participants’ chosen neutral posture. All images were taken with tablets (Galaxy Tab S7, Samsung, Suwon-si, South Korea) on a light blue card-stock photo background with yellow photo-scales placed distally from the hand. Some images also include a label (black text on white background) with an alphanumeric value that was used internally to map image files to other data from the same participant. A set of 49 points was manually placed on all images using ImageJ (Figure 2), and used to calculate 58 hand measurements previously defined by the United States 1988 Army Anthropometric Study.^9^ The conversion from pixels to centimeters was performed by placing the first two points one centimeter apart on the ruler included in all images and determining the number of pixels across that distance. Hand length and width were also measured directly from each participant’s hand using a ruler; these direct measurements served as independent verification of measurements derived from photogrammetry.

**Figure 2.**
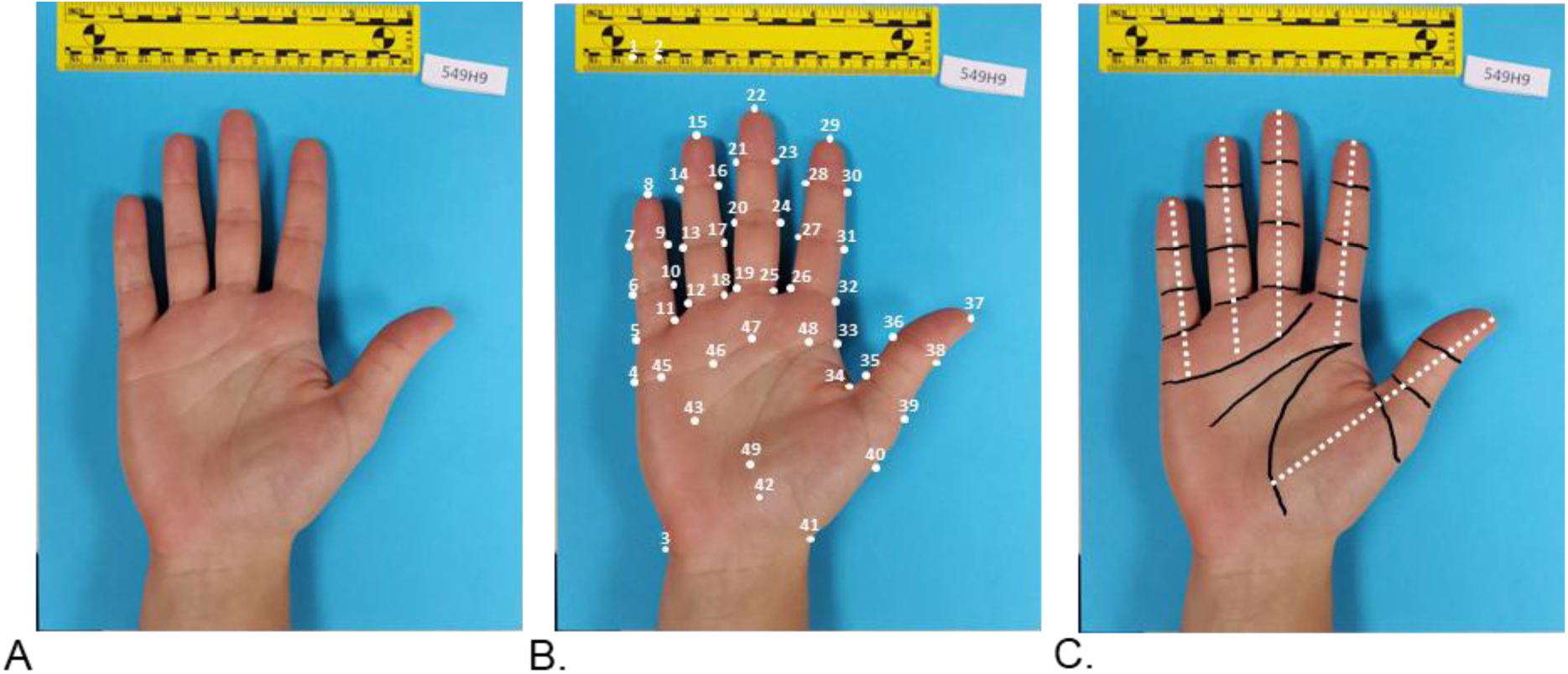
Example method of hand photogrammetry to determine hand measurements as defined by the United States 1988 Army Anthropometric Study. (a) Example of a hand image from the BHaM population dataset. Participants were instructed to place their hand flat on the blue background in a neutral finger position. The yellow ruler was included in all photos to provide scale and determine the pixel to centimeter ratio needed for photogrammetry. (b) A total of 49 points were placed on each hand image using ImageJ. The point set includes the key points defined by the United States 1988 Army Anthropometric Study as well as additional points within the palm. The additional points were included to refine the estimated link lengths. Originally, link lengths were determined by the palm crease lines defined by two straight lines (between points 4 & 47 and points 33 & 43). (c) The additional points (45, 46, 48, 42) were placed on the palm creases (black) to be in line with the digit axis (white dashes) which runs from the finger-tip along the midline of the finger joint creases (black).

### 2.2 BHaM Biomechanics Dataset

A subset of 30 participants [n=30; age 48.6 ± 23.9 years; 14 male, 16 female; 26 right hand dominant, 4 hand left dominant] were recruited to take part in a follow-up portion of the IRB-approved study (UF IRB #202002061), which used traditional biomechanics to understand the elbow, wrist, and hand. The first session evaluated elbow and wrist mechanics using surface electromyography (sEMG) and dynamometry. The second session evaluated hand mechanics using motion capture, fine-wire EMG, force recordings, and MR imaging from participants’ dominant arm.

Participants were originally recruited into the biomechanics dataset study to include 10 participants from three age ranges [young adults: 18-39 years, middle-aged adults: 40-63 years, and older adults: 64-95 years]. Five males and five females were to be recruited for each age group, and each set of five individuals was to be selectively recruited to represent the distribution of grip strength for their age/sex (Figure 3). This representative distribution was to be achieved by selecting (i) a participant whose grip strength was within 20 N of the age-sex mean, (ii) a participant within 20 N of the mean plus one standard deviation, (iii) a participant within 20 N of the mean minus one standard deviation, and (iv) a participant within 20 N of the mean plus two standard deviations, and (v) a participant within 20 N of the mean plus two standard deviations. However, due to recruitment constraints, 14 young adults [7 male, 7 female], 2 middle-aged adults [1 male, 1 female], and 14 older adults [6 male, 8 female] were recruited, while maintaining the criteria for representing a grip strength distribution of the total population (Figure 4).

**Figure 3.**
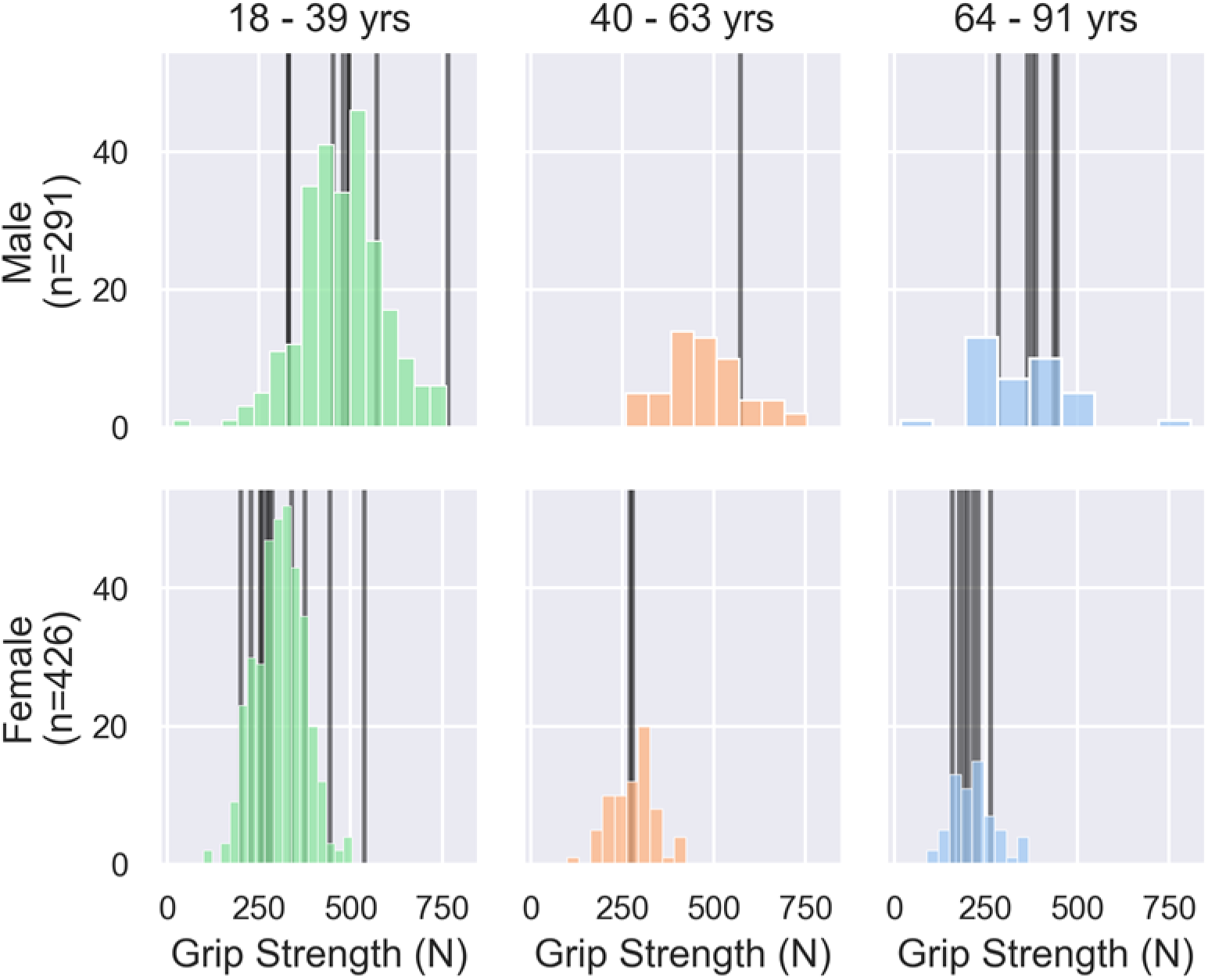
Grip strength distributions used to determine recruiting criteria for the biomechanics sample. Stratified grip distributions are shown in color with the grip strength of recruited participants shown by vertical black lines. The participants in the BHaM Biomechanics Dataset (black) were originally recruited by targeting an individual from the mean, ±1, and ±2 standard deviations for each sub-group. However, due to low recruitment rates from participants between 40-63 years (orange), and male participants over 64 years (blue), the recruitment criteria changed focus to capture the population of individuals under 39-years (green) and over 64 years (blue).

**Figure 4.**
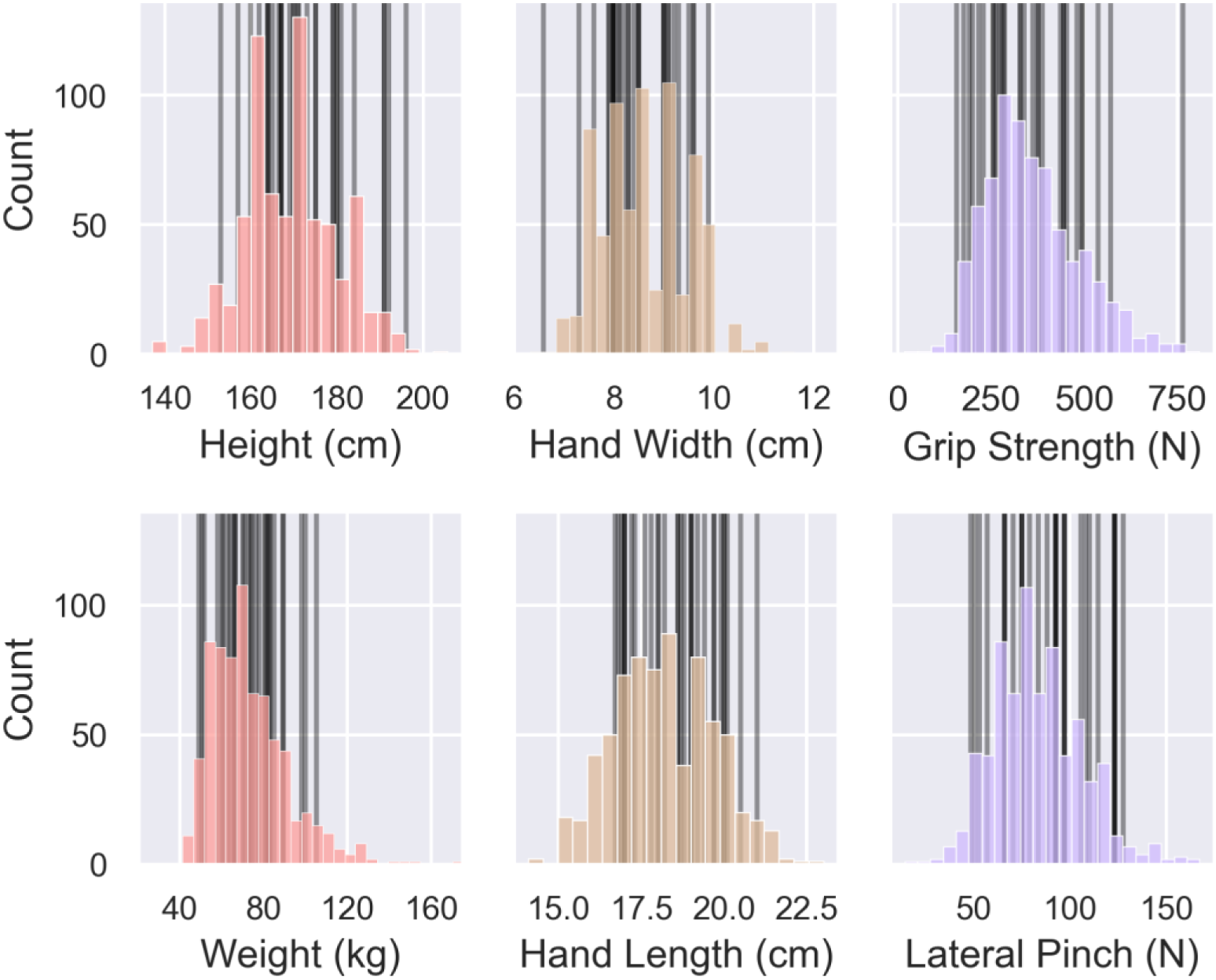
Distributions of all participants in the BHaM Population Dataset (colored) and BHaM Biomechanics Dataset (black lines) illustrating the population diversity captured within the biomechanics subset. Although recruitment for the BHaM Biomechanics Dataset was determined by selectively recruiting individuals based on grip strength, the selective recruitment still captured the distribution of other key metrics (height, weight, hand width, hand length, and lateral pinch). Recruited biomechanics participants included individuals from ±2 standard deviations for height, hand width, hand length, grip strength, and lateral pinch, and ±1 standard deviation for weight.

#### 2.2.1 Elbow and Wrist Session

Participants first completed the elbow and wrist testing session since it was noninvasive. Surface electrodes (Trigno Quattro Sensor and Trigno Avanti Sensor, Delsys, Massachusetts, United States) (3,000 Hz) were placed on the finger flexors [*flexor digitorium profundus, flexor digitorium superficialis*], the primary finger extensor [*extensor digitorium communis*], wrist flexors [*flexor carpi radialis*, *flexor carpi ulnaris*], wrist extensors [*extensor carpi radialis*, *extensor carpi ulnaris]*, biceps, and the triceps long head. Muscles were identified with palpation and confirmed with muscle-specific resistance^29^ Participants were then placed in a dynamometer to record analog joint angle, angular velocity, and torque (System Pro 4, Biodex, New York, United States).

Data acquisition of the sEMG and dynamometer systems were time synchronized via Vicon Nexus 2.14 (Lock Lab, Vicon, Oxford, England).

While seated in the dynamometer, each participant performed isometric maximum voluntary contractions (MVC) and 50% MVC for flexion and extension separately at the elbow and wrist. Participants also performed isokinetic full range of motion flexion/extension separately at the elbow and wrist. For isometric and range of motion wrist tasks, participants had the humerus of the dominant arm strapped to their thorax in a neutral position, the thorax double strapped to the chair back in a sitting position, elbow positioned at 90°, and the fully supinated forearm strapped to a forearm support to maximize wrist isolation (Figure 5). For isometric elbow tasks, the forearm support and strap were removed, but the elbow remained positioned at 90°. For elbow flexion/extension range of motion, the arm was elevated to be parallel to the floor and the only strap on the arm was positioned immediately proximal to the elbow and strapped to a support (Figure 5).

**Figure 5.**
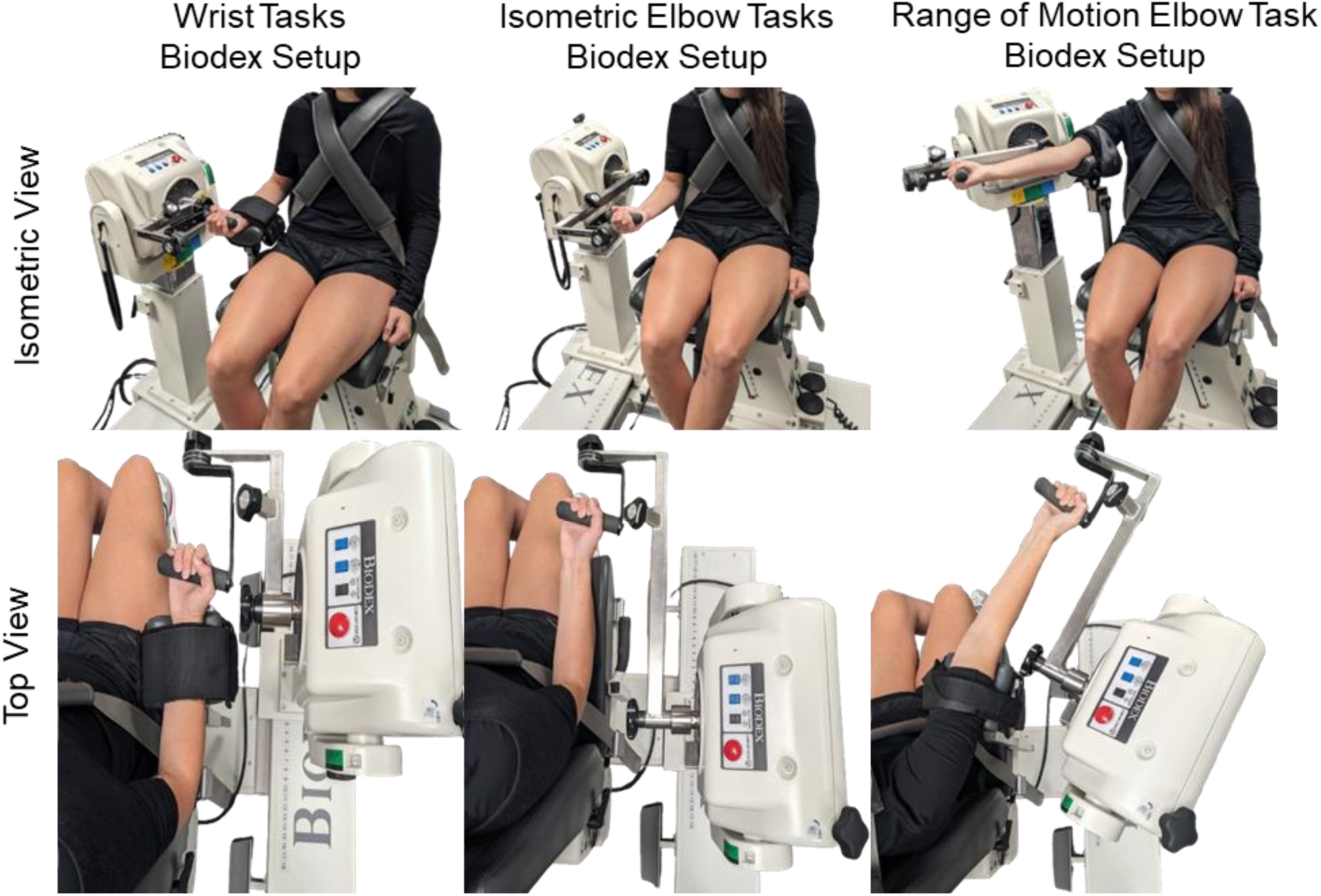
Dynamometer setup for wrist tasks, isometric elbow tasks, and isokinetic elbow tasks. Subjects were constrained to a Biodex System 4 Pro Chair using body straps to promote isolated movement of the targeted joint. During wrist tasks, the elbow was positioned at 90°, the wrist fully supinated, the forearm strapped to a support, and the center of the dynamometer positioned at the wrist joint center. During isometric elbow tasks, the elbow was positioned at 90°, the wrist fully supinated, and the center of the dynamometer positioned at the elbow joint center. During range of motion elbow tasks, the arm was elevated to be parallel to the ground and strapped to the arm support, the dynamometer arm was rotated to align with the forearm to account for the carry angle, and the center of the dynamometer was positioned at the elbow joint center.

During isometric tasks, participants performed five trials of 5 seconds with 10 second rest in-between. All isometric trials had real-time visual feedback in the form of a graph of the raw time-series z-component force signal. During 100% MVC tasks, participants were instructed to “get the signal as high up on the screen as possible,” while for 50% submaximal tasks. participants were instructed to “have the force as close as possible” to a horizontal target line placed on the graph at a level equivalent to 50% of the 100% MVC. During range of motion tasks, participants performed 5 cycles of full range of motion flexion-extension at 60 °/s, had a 30 second rest, then performed another 5 cycles at 120°/s. Task order within each task group (isometric, range of motion) were randomized. The elbow and wrist session took approximately 1.5 hours with 30 minutes for electrode placement and one hour for tasks.

#### 2.2.2 Hand Session

Baseline anthropometrics and strength measurements were taken prior to fine-wire EMG insertions to identify any changes caused by interference from sensors. A clinical baseline was established by using clinical dynamometers to assess hand strength. Grip strength was assessed with a digital Jamar grip dynamometer (DHD-1, B&L Engineering, California, United States); while lateral pinch, tip pinch, and isometric index flexion were assessed using a hand-held pinch gauge (10lb, 30lb, 60lb Pinch Gauges, B&L Engineering, California, United States). Since clinical dynamometry provides only one maximum value in a single force direction, we also assessed thumb-and finger-tip forces using a multi-degree-of-freedom, time-series force sensor. Time-series baselines were recorded for 5 MVC isometric tasks: jar grasp, lateral pinch, tip pinch, index flexion, and closing jar twist (Figure 6).This task set and set-up is similar to that originally described by <u>Halilaj et al., 2014</u>. Here, each task was performed for five 5-second trials with five seconds rest periods between tasks. Thumb-tip force during jar grasp, lateral pinch, and tip pinch as well as finger-tip force during index flexion were recorded with a six-degree-of-freedom force sensor (Mini40, ATI, North Carolina, United States) (3,000 Hz), while closing jar twist was record with a one-degree-of-freedom, time-series force sensor (Model D Load Cell, Honeywell, North Carolina, United States) (3,000Hz).

**Figure 6.**
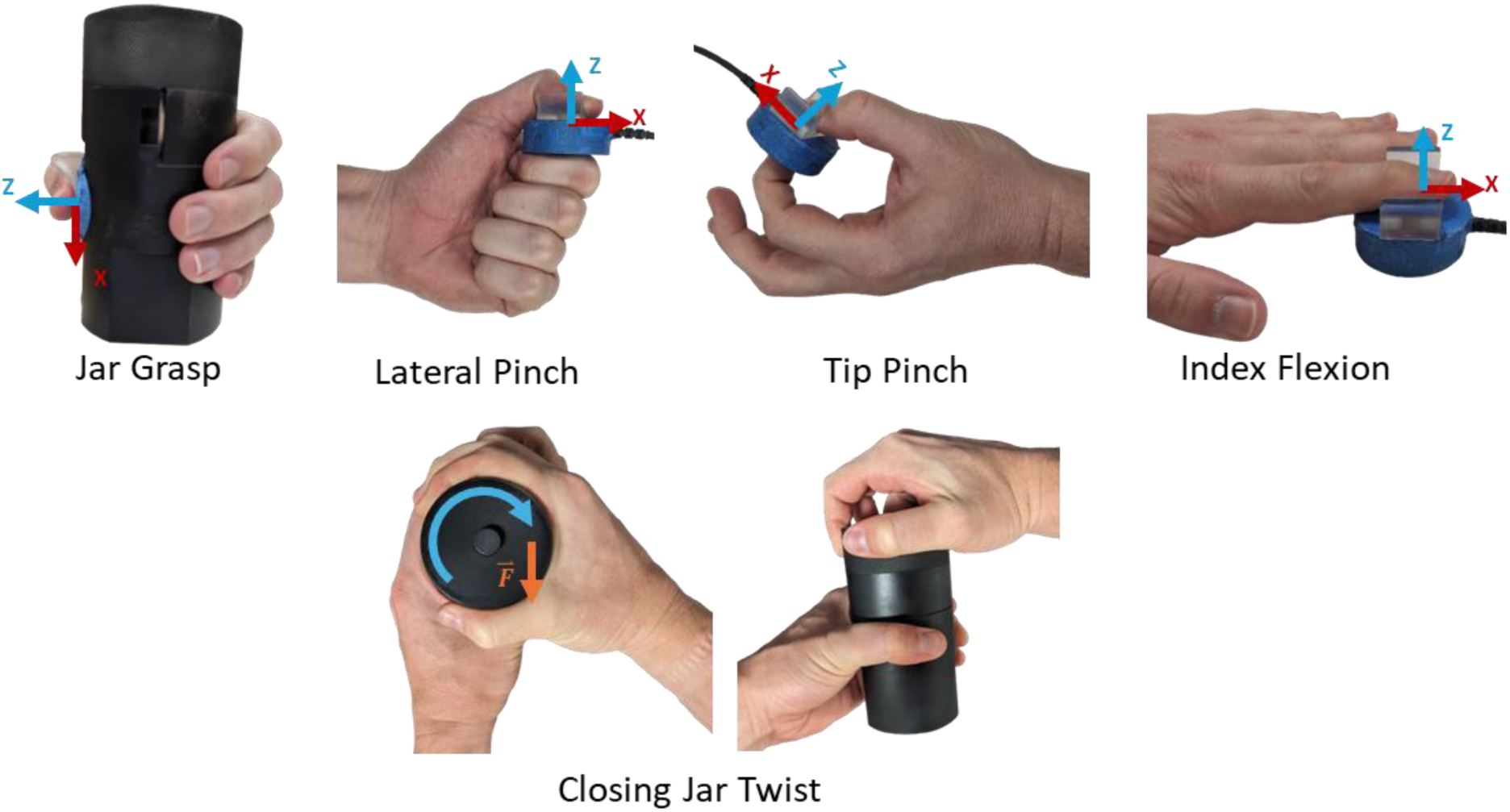
Five isometric hand tasks performed by participants with the 3D force coordinate system for each task. Tasks were chosen based on commonly performed daily tasks and previously published methods for recording thumb-tip force (Halilaj et al. 2014). For jar grasp, the six-degree-of-freedom force sensor was placed flush within a precut jar constrained to a table; the +z-axis was directed in the thumb’s dorsal direction and the +x-axis was directed in the direction of the table (radially). For lateral pinch, tip pinch, and index flexion the six-degree-of-freedom force sensor was aligned so that the +z-axis pointed in the dorsal direction and the +x-axis pointed in the distal direction. For jar twist, participants were only instructed to close the jar lid while a one-degree-of-freedom recorded the flossing force. Since participants self-selected their closing approach two primary twisting phenotypes were observed. The first observed closing method (left) was a cylindrical grasp, while the second observed closing method (right) was a spherical grasp.

Following baseline force measurements, fine-wire electrodes (30mm x 27g single-use paired fine wire electrode, Rhythmlink, South Carolina, United States) were used to record muscle activity from 8 thumb muscles (Trigno Avanti Spring Contact Adapter, Delsys, Massachusetts, United States) (3,000 Hz). Electrodes were placed under ultrasound guidance (SuperSonic Imagine Mach 30, SuperSonic Imagine, Provence, France) in four extrinsic [*flexor pollicis longus* (FPL), *extensor pollicis longus* (EPL), *extensor pollicis brevis* (EPB), and *abductor pollicis longus* (APL)] and four intrinsic [*flexor pollicis brevis* (FPB), *adductor pollicis* (ADD), *opponens pollicis* (OPP), and *first dorsal interossei* (FDI)] muscles.^29^

Motion capture was recorded with a 12-camera motion capture system (Vicon Vero, Vicon, Oxford, England) (100 Hz) and a custom 44-count marker set (Figure 7). The thumb was tracked with six markers on the first metacarpal [three proximal head, three distal head], three markers on the proximal phalanx, and three markers on the distal phalanx. The fingers were tracked with a single marker on the distal metacarpal head, proximal, medial, and distal phalanx. Tracking clusters were placed on the hand, forearm, upper arm, and thorax [acromion, sternal notch, C7, T12]. Additional scaling makers were placed on the radial styloid, ulnar styloid, medial elbow, and lateral elbow.

**Figure 7.**
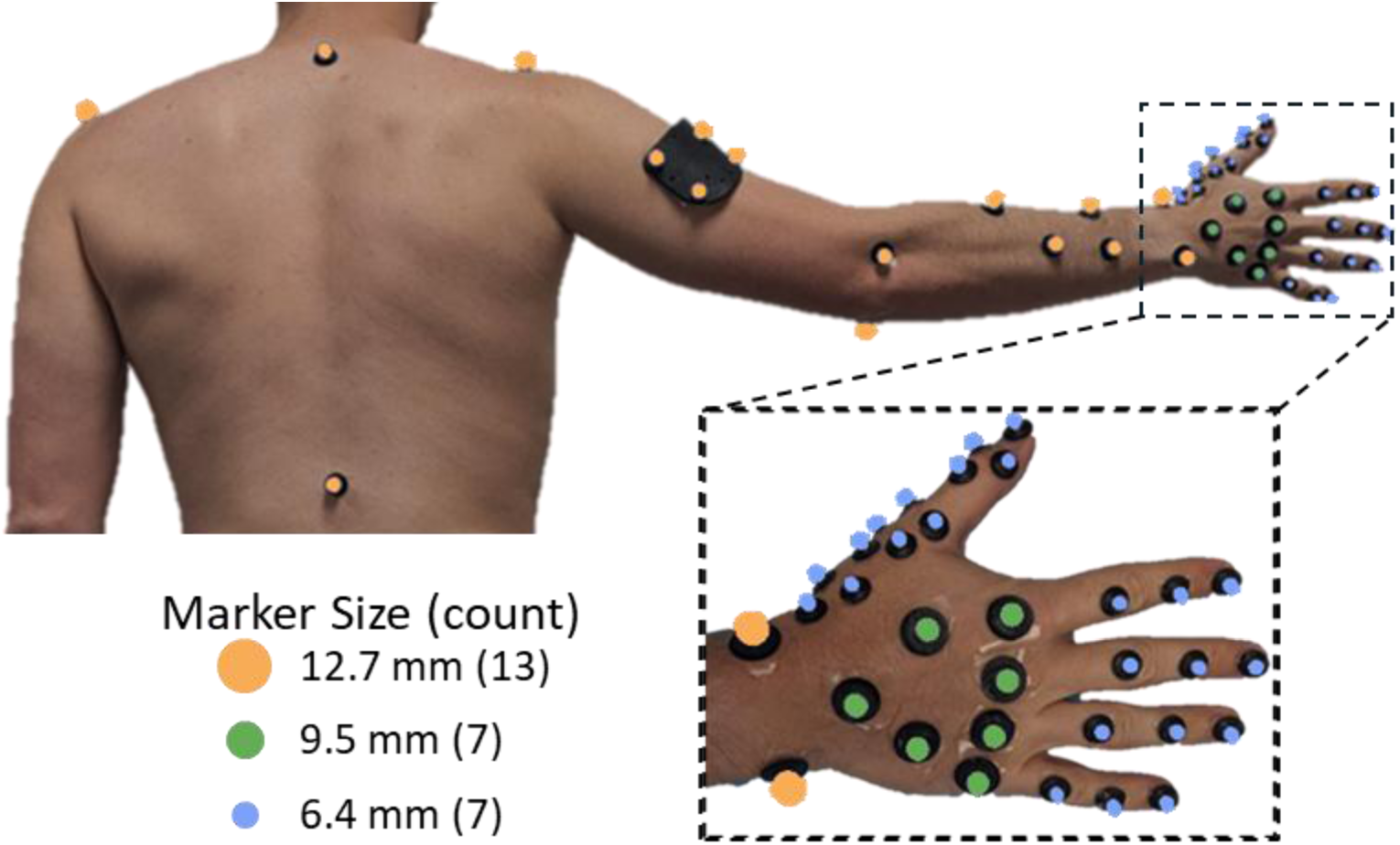
Posterior view of the custom 44-count motion capture marker set (jugular notch marker not shown) used to track the thorax, arm, forearm, hand, thumb, and digits. The forearm cluster varied between participants to accommodate wires and boxes (not shown) required for fine wire EMG. Three sizes of retroreflective markers are used (12.7mm: orange, 9.5mm: green, 6.4mm: blue) to accommodate the marker density on the back of the hand and prevent large markers from interfering with participants’ movements.

To enable EMG normalization, muscle specific MVC were performed five times for 5 seconds with 5 second rest periods, while a trained experimentalist provided defined resistance for each muscle.^29,30^ To facilitate motion capture data processing, a static pose was taken with participants seated with their arm extended out in front of them [shoulder flexion: 90°, elbow flexion: 0°, wrist supination/deviation: 0°].

Each participant performed seven range of motion tasks and five isometric tasks to capture hand and thumb functions. The seven range of motion tasks (Figure 8) were each performed 5 times to a metronome beat (60 bpm). The five isometric tasks (Figure 6) were then performed 5 times at both 100% and 50% MVC with each trial involving exerting for 5 seconds with 5 second rest between trials. Similar to the baseline measurements, thumb-and finger-tip force was measured in four tasks (lateral pinch, tip pinch, jar grasp, index flexion) with a six-degree-of-freedom, time-series force sensor (Mini40, ATI, North Carolina, United States) (3,000Hz), while a one-degree-of-freedom, time-series force sensor (Model D Load Cell, Honeywell, North Carolina, United States) was used to record thumb-tip force during closing jar twist (3,000Hz). During jar twist, participants were instructed to “close the jar” how they normally would. This instruction resulted in two common methods, either (i) using only the fingertips or (ii) wrapping the fingers around the lid. Task order within each task group (baseline, isometric) were randomized. Experimental procedures took approximately two hours with one hour for baseline measurements and fine-wire insertions, and one hour for marker placement and completing tasks.

**Figure 8.**
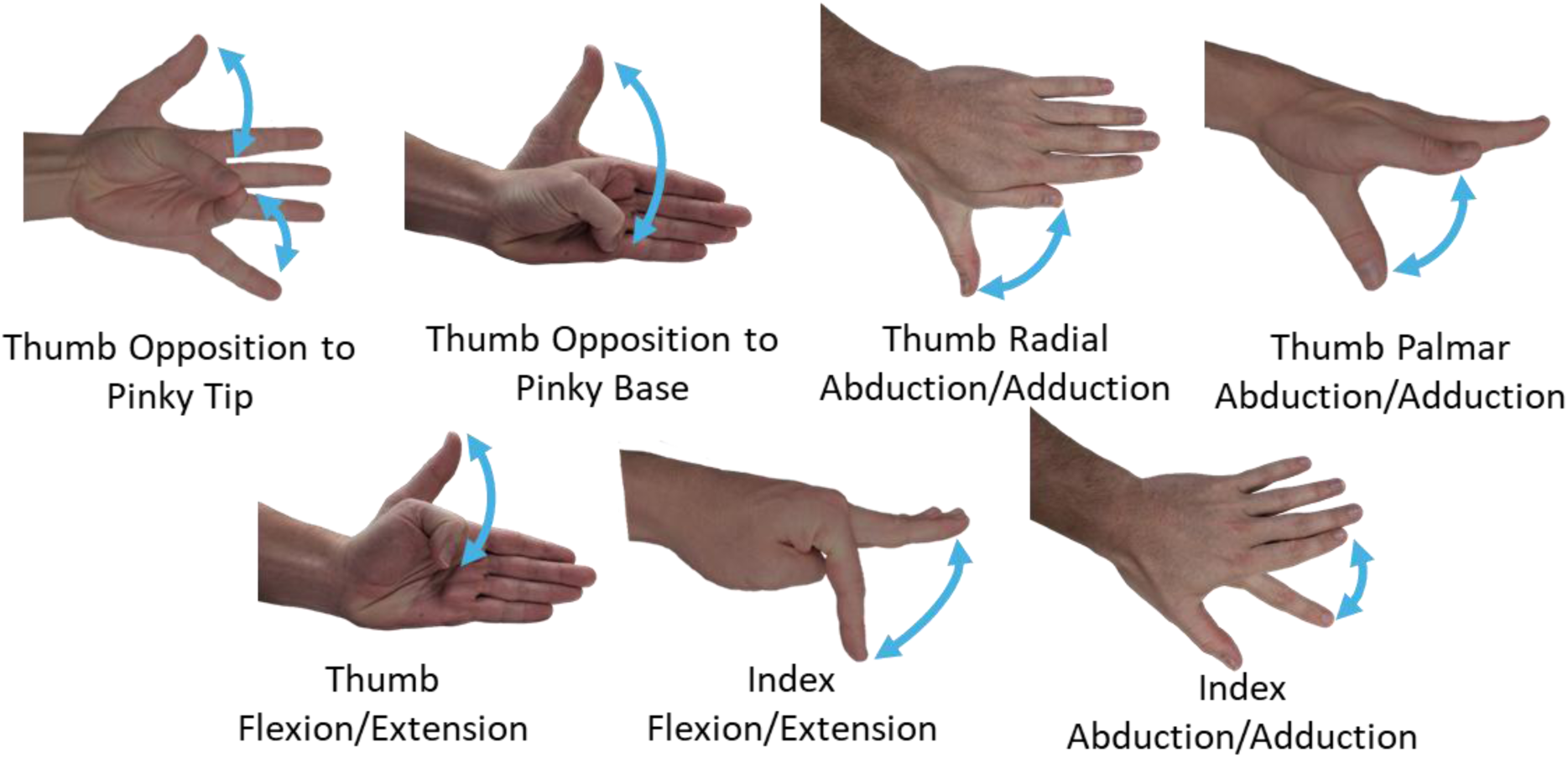
Range of motion tasks captured using optical motion capture. Tasks were chosen to capture the primary motions of the thumb and second metacarpophalangeal joint.

#### 2.2.3 MR Imaging Session

A subset of 15 participants [n=15; age: 46.1 ± 23.3 years; 7 male, 8 female; 12 right hand dominant, 3 left hand dominant] were eligible for structural MR imaging of the arm and forearm taken two weeks after the biomechanics sessions (UF IRB #202202865). Eligibility was determined by 1) MR imaging exclusion criteria (e.g., ferrous implants, claustrophobia, etc.),^31^ 2) quality of experimental data from the biomechanics sessions, and 3) maintaining demographic representation. Participants were placed supine into the scanner while 3T axial MR images were acquired (ST: 3mm, TR: 3,440 ms, TE: 20 ms, FA: 90°, FOV: 11cm, matrix: 224 x 224, bandwidth: ±1 kHz) (3 T/70 cm Philips MR7700, Philips, Amsterdam, Netherlands).^20,32^ Forearm muscles and bones (Table 1) were manually segmented in 3D Slicer by a single, trained researcher.^33^ Segmentations were then summarized by 3D Slicer’s volume measurement tool. Segmentations were evaluated for consistency by ensuring segmented muscle volumes fell within two standard deviations of previously published distributions from previous forearm MR volume studies.^20,32^

**Table 1.** Segmented forearm muscles and bones from MR images.

| Name | Abbreviation |
| --- | --- |
| *Abductor pollicis longus* | APL |
| *Anconeus* | ANC |
| *Brachioradialis* | BRD |
| *Extensor carpi radialis* | ECR |
| *Extensor carpi ulnaris* | ECU |
| *Extensor digiti minimi* | EDM |
| *Extensor digitorum communis* | EDC |
| *Extensor pollicis brevis* | EPB |
| *Extensor pollicis longus* | EPL |
| *Flexor carpi radialis* | FCR |
| *Flexor carpi ulnaris* | FCU |
| *Flexor digitorum profundus* | FDP |
| *Flexor digitorum superficialis* | FDS |
| *Flexor pollicis longus* | FPL |
| *Palmaris longus* | PL |
| *Pronator teres* | PT |
| *Supinator* | SUP |
| Radius | RAD |
| Ulna | ULN |
| Adipose tissue | FAT |

### 2.3 Data Processing

Kinetic data was low-pass filtered to remove high-frequency noise. The six-degree-of-freedom forces were low-pass filtered (4 Hz,4^th^ order-Butterworth) and rotated 22.5° about the Z-axis (dorsal-ventral). The rotation was performed to align the sensor and experimental reference frames (Figure 6). This alignment was necessary as the sensor cable was used to align the sensor with the proximal-distal direction of the participant’s arm. In a similar manner, the jar twist force was filtered with a 4 Hz, 4^th^ order-Butterworth low-pass filter. The elbow and wrist session’s analog signals (position, velocity, torque) were low-pass filtered (0.5 Hz, 4^th^ order-Butterworth) to match the dynamometer’s auto-generated torque (System Pro 4, Biodex, New York, United States).

EMG signals were processed to obtain normalized activation envelopes. Signals were high-pass filtered (20 Hz, 4^th^ order-Butterworth) to remove motion artifacts. Signals were then rectified to remove negative voltages. Finally, EMG envelopes were then obtained with a low-pass filter (5 Hz, 4^th^ order-Butterworth) and smoothed with a 0.2 s moving average filter. All EMG signals were normalized using the maximum value recorded across all isometric and muscle-specific MVC tasks.

Marker trajectories were processed in preparation for inverse kinematics. Trajectories were manually corrected for marker toggling and gap-filled in Vicon Nexus 2.14 (Vicon, Oxford, England). Gap-filled trajectories were then low-pass filtered to remove high frequency motion artifacts and marker jitter (6 Hz, 4^th^ order Butterworth). Markers were rotated 180° about the y-axis (superior-inferior) to align the experimental coordinate frame with the OpenSim global coordinate frame. Trajectories from left-handed participants were mirrored about the sagittal plane to simulate a right arm.

### 2.4 Calculating Joint Angles During Tasks

A generic musculoskeletal model was anthropometrically scaled to each participant’s height and weight for inverse kinematics. The generic model was created by combining the MoBL-ARMS upper limb^6,8^ and ARMS hand and wrist models^5,7^ and adding the custom marker set (Figure 7). In the generic model, for scaling purposes, the thorax mass was changed from 0 kg to 20.46 kg based on the Thoracoscapular shoulder model.^34^ The generic model was then scaled using the maker data from the static pose. During scaling, since the model consisted of only the thorax and right arm, model mass was set to 34.17% or 38.90% of total mass for female and male participants, respectively.^35^

Joint angles were calculated using inverse kinematics performed in OpenSim 4.4 for isometric and range of motion hand tasks.^36^ Inverse kinematics was performed with an iterative batch processing script that altered marker weights on the thumb or index finger between 1, 10, and 100 while adjusting the start time by 0.01, 0.02, or 0.1 s. Batch processing results were then verified in the OpenSim visualizer and the weights and start times of failed simulations were manually adjusted until a solution was found. This iterative approach was required because variations in marker weights and start times can cause the OpenSim inverse kinematics tool to either i) fail to run to completion for a given trial or ii) generate physically impossible results.

### 2.5 Defining Task Onset-Offset Timing

The start and stop of each task trial (i.e., trial bands) were identified with a Gaussian mixture model, which is an unsupervised method for data clustering.^37^ The Gaussian mixture model assumes data is generated from a finite number of normal distributions. The model then estimates the number of normal distributions needed to represent the data. Data points are then clustered based on the distribution from which each data point has the highest probability of belonging. When implementing a purely unsupervised approach, the number of clusters is determined by the Gaussian mixture model; however, the number of clusters can be constrained leading the Gaussian mixture model to only identify the shape of a prescribed number of distributions. Methods using Gaussian mixture models have been found to be better for batch processing data compared to thresholding methods because Gaussian mixture models are capable of handling variance in baseline noise, duration (or length), and measurement modality.^38,39^ Here, the Gaussian mixture model was set to identify two clusters (during trial vs. during rest period), thereby creating a binary label for each signal. The binary label was then eroded and dilated by 25% of trial length to remove mislabeled regions. Then the label was dilated and eroded by 25% of trial length to fill gaps within the trial regions. For elbow and wrist trials, the measured torque was used to identify trial bands of isometric tasks, and joint angle was used to identify trial bands for range of motion tasks. For hand trials, the dorsal-ventral force component was used to identify trial bands for isometric force tasks, and the calculated joint angles were used to identify trial bands for range of motion tasks. For EMG MVC trials, the EMG signal for the targeted muscle was used to identify trial bands.

## 3 Data Records

The BHaM database is publicly available at Kaggle (https://doi.org/10.34740/kaggle/ds/7895310) and the files are summarized in Table 2.^40^

**Table 2.**
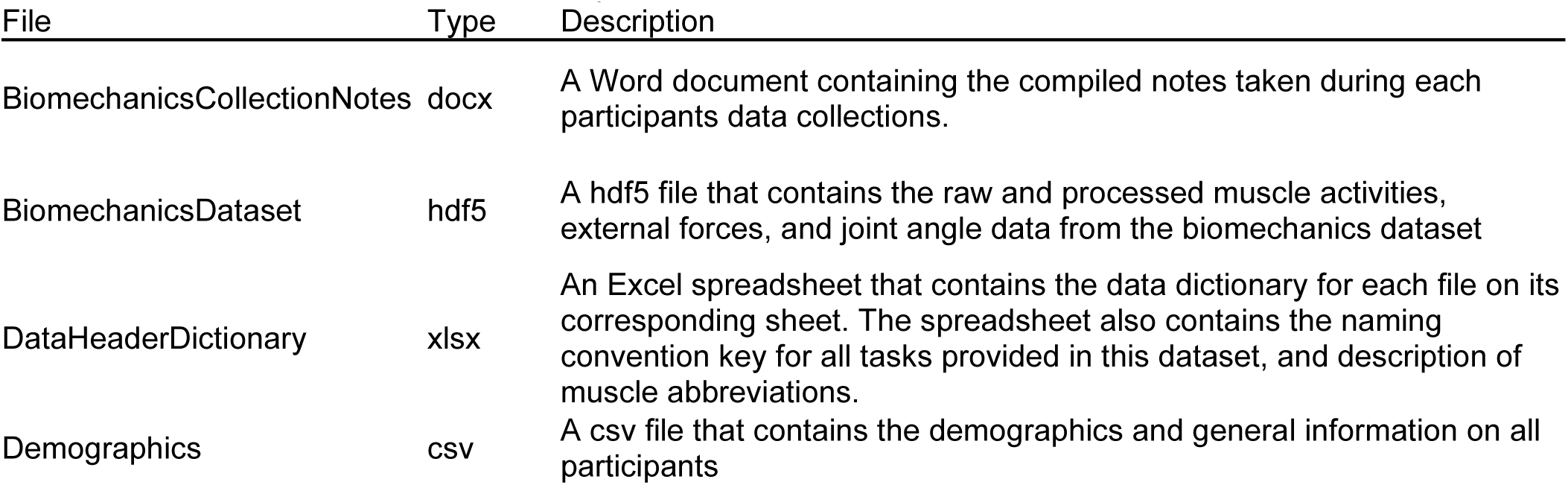
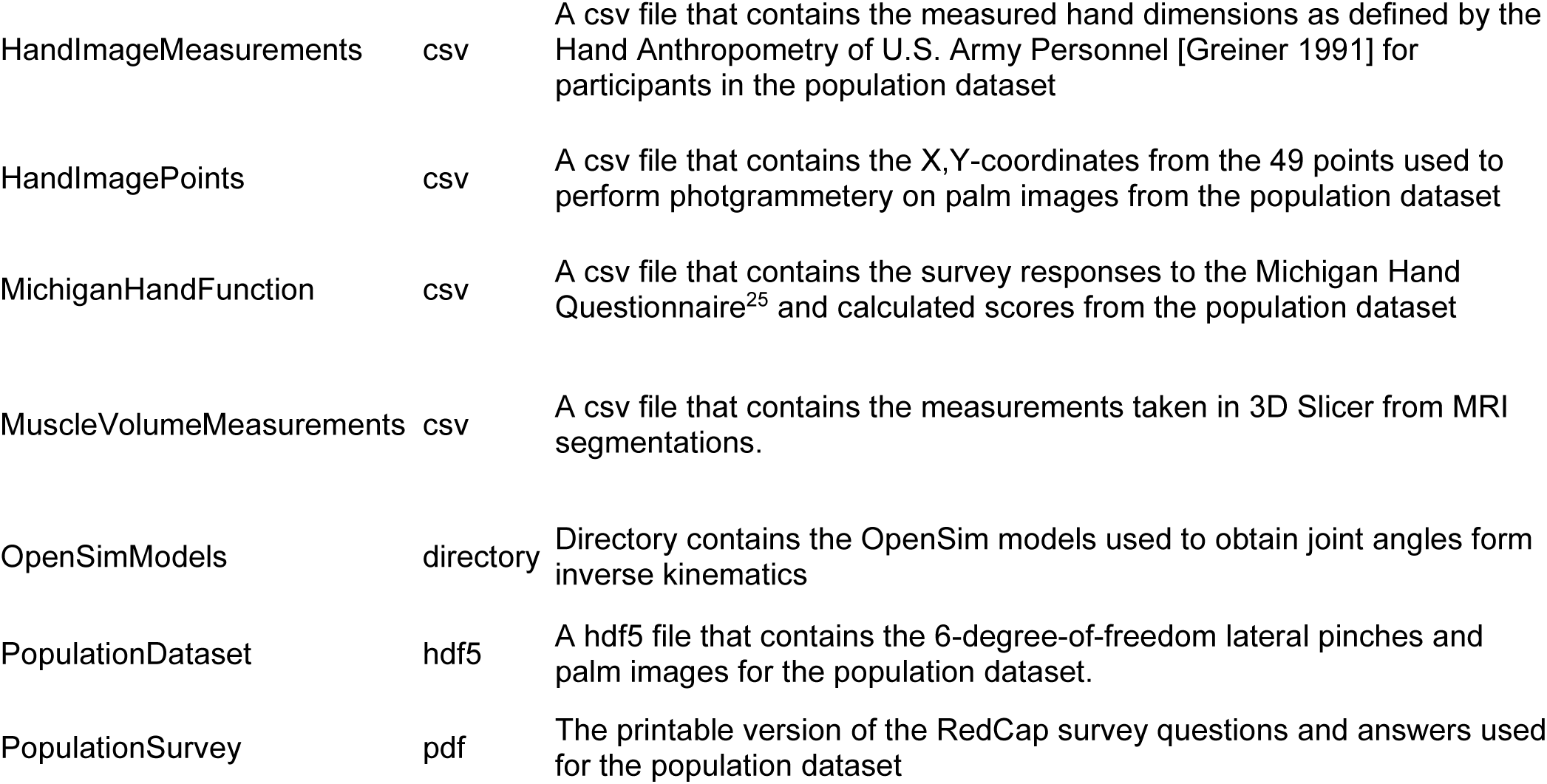
Published dataset files and descriptions.

| File | Type | Description |
| --- | --- | --- |
| BiomechanicsCollectionNotes | docx | A Word document containing the compiled notes taken during each participants data collections. |
| BiomechanicsDataset | hdf5 | A hdf5 file that contains the raw and processed muscle activities, external forces, and joint angle data from the biomechanics dataset |
| DataHeaderDictionary | xlsx | An Excel spreadsheet that contains the data dictionary for each file on its corresponding sheet. The spreadsheet also contains the naming convention key for all tasks provided in this dataset, and description of muscle abbreviations. |
| Demographics | csv | A csv file that contains the demographics and general information on all participants |
| HandImageMeasurements | csv | A csv file that contains the measured hand dimensions as defined by the Hand Anthropometry of U.S. Army Personnel [Greiner 1991] for participants in the population dataset |
| HandImagePoints | csv | A csv file that contains the X,Y-coordinates from the 49 points used to perform photogrammetry on palm images from the population dataset |
| MichiganHandFunction | csv | A csv file that contains the survey responses to the Michigan Hand Questionnaire<sup>25</sup> and calculated scores from the population dataset |
| MuscleVolumeMeasurements | csv | A csv file that contains the measurements taken in 3D Slicer from MRI segmentations. |
| OpenSimModels | directory | Directory contains the OpenSim models used to obtain joint angles from inverse kinematics |
| PopulationDataset | hdf5 | A hdf5 file that contains the 6-degree-of-freedom lateral pinches and palm images for the population dataset. |
| PopulationSurvey | pdf | The printable version of the RedCap survey questions and answers used for the population dataset |

Brief description of the provided data files is provided below:

- *Kinematics, Kinetics, & Muscle Activity:* The **signal data (time series forces, EMG, joint angles, hand images)** for both the population and biomechanics datasets are stored as Hierarchical Data Format version 5 (HDF5) files. Each HDF5 file includes the participant identifier as the top-level group with each participant’s demographics included within the group metadata.
- *Hand Images & Anthropometrics:* The **hand anthropometric data** is provided in two forms. First, the coordinates of the 49 hand points and the 58 hand measurements are provided for all participants as two comma separated files (csv) (HandImageMeasurements.csv, HandImagePoints.csv). Second, participant-specific values are embedded within the HDF5 metadata for the published hand images. Because the informed consent process involved the opportunity for participants to choose to have (or not have) imaging data publicly shared, the published hand images are limited to those for which participants provided consent to share, and did not contain identifying markings (e.g., tattoos). In the published dataset, 509 of the 726 hand images are fully available.
- *Demographics & Self-Reported Hand Function:* The population dataset’s **survey responses** (Demographics.csv, MichiganHandFunction.csv) are stored as comma separated files (csv). A printed version of the **RedCap survey** (PopulationSurvey.pdf) is provided to show the phrasing of the questions and survey response choices.
- *MR Imaging Data:* The **muscle segmentation measurements** (MuscleVolumeMeasurents.csv) are stored as comma separated files (csv). The original DICOM files and segmentations are not published due to data privacy requirements, but these files are available upon request from the study team.
- *Descriptors:* The **data descriptors**, **task descriptors**, and **muscle abbreviations** for all published data are listed within DataHeaderDictionary.xlsx with a separate sheet for each file.
- *Experimental Notes:* The **participant notes** from the biomechanics dataset experimental sessions were edited for conciseness and clarity (BiomechanicsCollectionNotes.pdf). The participant notes consist of notes taken by the experimenter during data collections to document unexpected events (equipment failures, experimenter errors, participant comments, abnormal signals, etc.) that occurred throughout a participant’s experimental session.
- *OpenSim Modeling & Simulations:* The **generic OpenSim model** (GenericModel.osim), **participant scaling settings**, and **participant scaled models** are included in a directory for all OpenSim models (OpenSimModels). Note, use of the models is limited by the licenses of the original models (MoBL-ARMS upper extremity; ARMS hand and wrist) from which the BHaM models are derived; licensing information is included in each model file.

## 4 Technical Validation

For technical validation, we assessed whether the BHaM database (i) confirmed known differences between pain and demographic characteristics, (ii) confirmed known differences between pain and hand function, and (iii) identified new knowledge regarding the differences between pain and hobby participation. These analyses used the following data from the population dataset:

- ***Pain Sub-score from the Michigan Hand Questionnaire***^25^ *(Pain (score) from MichiganHandFunction.csv)*: The pain sub-score is a self-reported pain metric based on five questions about the dominant hand. Each question is scored on a 1 to 5 scale, which is then averaged and transformed to a 0 to 100 scale. Higher numbers indicate worse pain.
- *Difficulty performing Activities of Daily Living (ADLs) from the Michigan Hand Questionnaire*^25^ *(12 ADLs from MichiganHandFunction.csv)*: The 12 ADL-related questions of the Michigan Hand Questionnaire use a 1 to 5 scale (“not at all difficult” to “very difficult”) to assess the participant’s ability to use their hands to perform ADLs. Five questions focused on ADLs performed using the dominant hand and seven questions focused on ADLs performed using both hands.
- ***Demographics*** *(Age, Sex, and Race_Ethnicity from Demographics.csv)*: Age (in years), biological sex (male, female, intersex, prefer not to answer) and race/ethnicity (Asian; Black, African American, or African; Hispanic, Latino, or Spanish; Middle Eastern or North African; Native American; or White) were self-reported.
- ***Hand Dominant Hobbies or Activities*** *(from Demographics.csv)*: An open-ended free response questions, which is independent of the Michigan Hand Questionnaire, was included in the population survey. The question read “List any hobbies or activities you have previously or are currently participating in that require you to train your hand/wrist (tennis, guitar, video games, basketball, sculpting, etc.).”

### 4.1 Stratification for Statistical Analyses

The full population dataset (n = 726 participants) was first cleaned to remove participants with missing data. A total of 10 participants were removed from the analysis due to missing pain scores, and 1 participant was removed due to an incomplete age field. This resulted in a total of 715 participants available for analysis.

The cleaned population dataset (n = 715 participants) was then stratified based on participant demographics to create three different analysis datasets (Figure 9). Stratification was performed based on (i) age, (ii) race/ethnicity, or (iii) biological sex. The age dataset included 715 participants stratified into three groups: young (18-39 years), middle-aged (40-63 years), and older (64-95 years) adults. The age cutoffs reflect previously established stratifications.^41,42^ The race/ethnicity dataset included 627 participants from four racial/ethnic groups: Asian, Black, Hispanic/Latino, or White. Although the population dataset includes participants from six racial/ethnic groups, our analysis focused on the four largest to ensure statistical power. Participants who identified themselves as multiracial (n = 69) were excluded from the race/ethnicity analysis to maintain focus on distinct racial/ethnic groups. The biological sex dataset included 707 participants, who identified as male or female. Participants who responded as Intersex (n = 3) or Prefer Not to Answer (n = 5) were excluded from analysis due to insufficient sample sizes for analysis.

**Figure 9.**
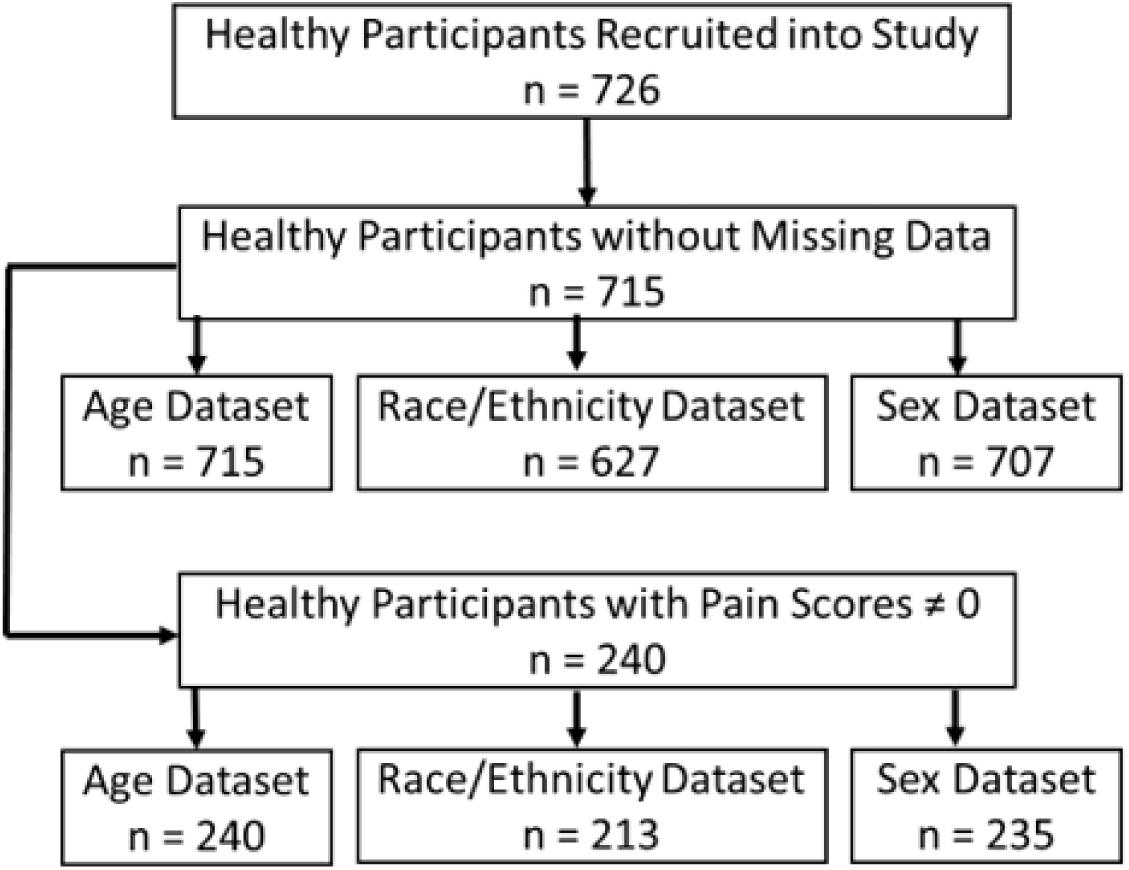
Flowchart illustrating the definition of the age, race/ethnicity and biological sex datasets. The demographic datasets were further stratified by pain score (primary analysis on all participants; sensitivity analysis on participants with pain score greater than zero) and then hobby (hand dominant vs. no hand dominant hobbies).

Each demographic analysis dataset was further stratified into two groups: individuals with hand-dominant hobbies and individuals without hand-dominant hobbies. This stratification enabled analysis of how self-reported engagement in hand-dominant hobbies influences self-reported pain scores. Individuals with hand-dominant hobbies were manually identified based on the free response data. The binary classification of individuals with versus without hand-dominant hobbies was then reviewed to ensure that individuals were not mislabeled. This review led to four individuals being moved from the hand-dominant hobbies to the no hand dominant hobbies group.

Finally, to evaluate the robustness of the reported results, a sensitivity analysis was performed that included only participants with non-zero pain scores. This subset allowed the opportunity to confirm known relationships, such as the relationship between pain and functional limitations.^43,44^ This subset also allowed the opportunity to explore understudied relationships such as pain across various demographic groups.^45,46^ The reproducibility of expected findings within this sensitivity analysis supports the validity of the population dataset and our analytical approach. The sensitivity analysis also elucidated whether there were floor effects that may have influenced observed group differences.

In the end, this means that all stratified analyses were conducted twice; first including all participants (termed the primary analysis dataset), and then excluding participants with a pain score of zero (termed the sensitivity analysis subset). These analyses were further stratified by hobby (hand dominant vs. no hand dominant hobbies), and comparisons were made across race/ethnicity, biological sex, and age.

### 4.2 Pain Across Demographic Groups

Statistical analyses to assess whether hand pain differed across demographic groups were performed in R (RStudio 2024.04.2+764). The normality of the pain score distributions across all demographic groups was assessed using the Shapiro-Wilk test. In all cases, the data was not normally distributed (p < 0.05) motivating the use of non-parametric statistical tests.

The average reported pain score across all groups was 13.8 (range 0 to 100). Although the population dataset was intended to reflect a healthy adult population, the pain data indicates that a subset of the sampled population may have had underlying hand or upper extremity disorders that caused pain. Pain data (Figure 10) generally aligned with known demographic trends, with mean pain scores being higher in females versus males,^47^ older adults versus younger adults,^48–50^ and black participants versus white participants.^51,52^

**Figure 10.**
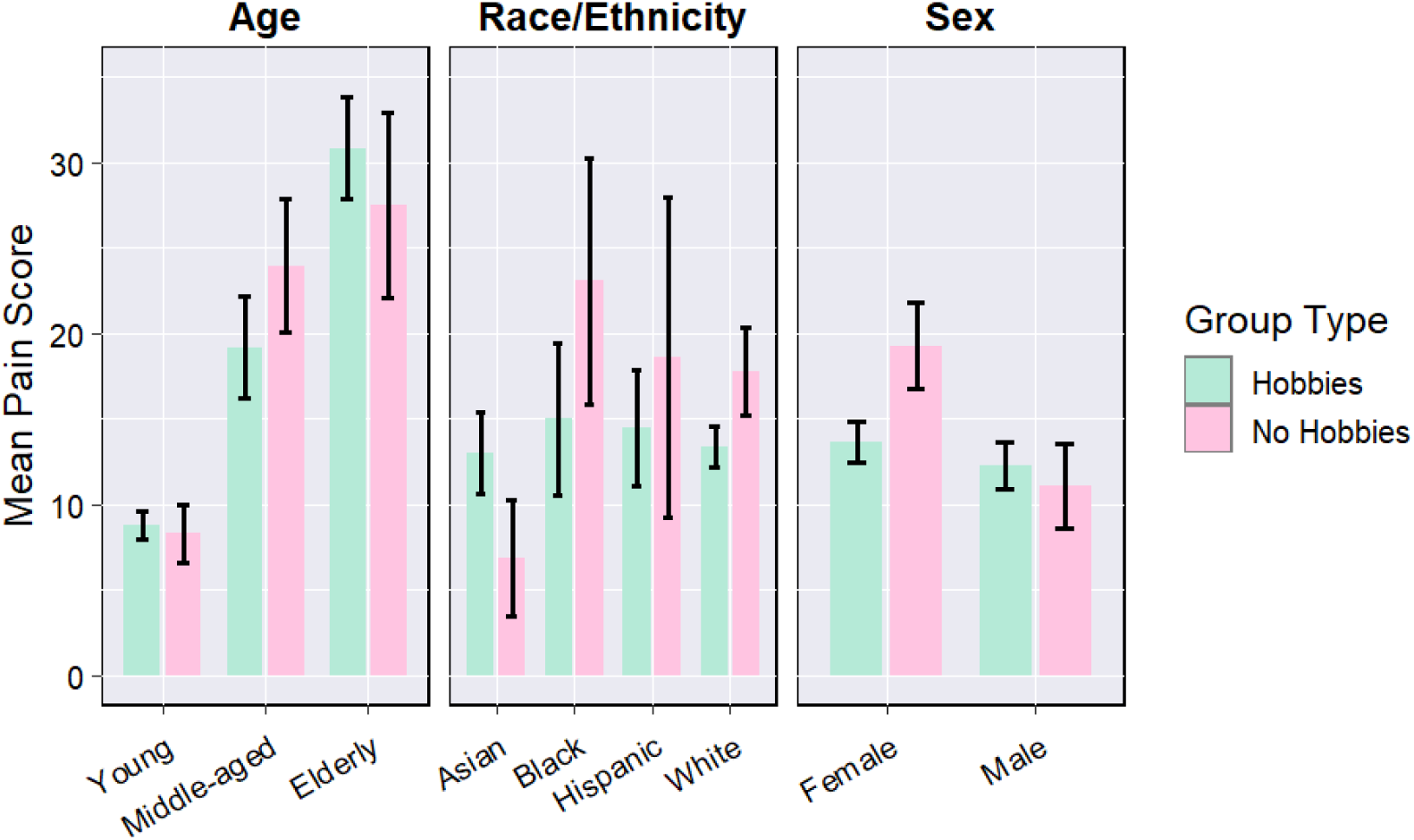
Mean pain scores for participants with (green) and without (pink) hand-dominant hobbies across three demographic groups. Error bars represent standard error.

### 4.3 Relationship between Pain and Function Across Demographic Groups

To assess the differences between pain and function, the 12 ADLs were analyzed. Specifically, Kruskal-Wallis tests were used to examine whether the reported mean pain score differed across the five levels of difficulty for performing an ADL. Differences were evaluated for each ADL and each demographic group. To correct for multiple comparisons, and reduce the likelihood of type I errors, the Benjamini-Hochberg false discovery rate correction test was applied.

When examining age, young adults did not exhibit as many significant differences post-correction as middle-aged and older adults; this aligns with prior work indicating that young adults have greater functional capabilities compared to older age groups.^53^ In contrast, middle-aged adults demonstrated significant pain-ADL differences across all ADLs, supporting prior work that indicates the onset of functional limitations and the difficulty with carrying out ADLs experienced by middle-aged adults.^54,55^ This may reflect a transitional timepoint where the onset of musculoskeletal conditions, such as hand osteoarthritis, is beginning to impact daily hand function.^56,57^ Older adults demonstrated significant pain-ADL differences across most ADLs; this is expected due to age-related functional decline.^43^

When examining race/ethnicity, differences between pain and difficulty performing ADLs were statistically significant (p<0.05) for all 12 ADLs in White participants. In contrast, significant pain-ADL differences were identified in 10, 5, and 1 ADL(s) for the Black, Hispanic, and Asian groups, respectively. This contrasts with a prior research study indicating that minority groups such as Black and Hispanic participants often report higher pain intensity than White participants, and Black participants solely reported higher pain interference in ADLs when compared to White participants.^58^ It is important to note that the BHaM population dataset is predominantly White, which can limit the generalizability of analyses to more diverse populations.

When examining biological sex, female participants demonstrated significant differences between pain and function for all ADLs, while male participants demonstrated significant difference for all ADLs, except holding a glass of water, eating using utensils, and washing hair. These patterns within our study may reflect broader sex differences of how pain is perceived and the risk of ADL limitations across biological sex. Prior research has reported that women report greater pain frequency,^59^ which can impact functional outcomes. Furthermore, women tend to have a higher overall risk of ADL limitations,^60^ which may be a disparity exacerbated by the increased prevalence of pain in women.

Overall, the pain-ADL differences identified in the BHaM population dataset align with the findings of previous research linking pain to functional limitations.^61^ Within our study, difficulty with jar opening, frying pan holding, and picking up a coin had significant differences with pain scores across almost all demographic groups. The pain-ADL relationships were most evident in White, female, and older adult groups.

### 4.4 Relationship between Pain and Hand-Dominant Hobbies

Statistical analyses to assess whether hand pain differed between participants with and without hand-dominant hobbies were also performed in R (RStudio 2024.04.2+764). Specifically, the Kruskal-Wallis’ test was used to evaluate group differences in pain score across demographics for both the hand-dominant hobby and no hand-dominant hobby subsets (Figure 10). Post-hoc testing was then performed by a pairwise Wilcoxon rank-sum tests with Bonferroni adjustments when appropriate.

Among participants with no hand-dominant hobbies, there were notable differences between the analysis that included all pain scores versus only non-zero pain scores (Figure 11). These differences suggest that the observed group differences in the primary analysis dataset may be influenced by individuals who report no pain, highlighting the importance of accounting for floor effects in pain-related research analyses using the BHaM database. In the primary analysis dataset, young participants reported substantially lower pain scores compared to middle-aged and older participants. However, in the sensitivity analysis subset, the difference in pain scores by age group decreased noticeably, as young participants’ average pain scores rose while the average pain scores for middle-aged and older participants remained stable. Similarly, when evaluating race/ethnicity and sex in the primary analysis dataset, Asians and males reported lower pain scores; but these differences were less evident in the sensitivity analysis subset, where we could see that the pain levels elevated for these two groups.

**Figure 11.**
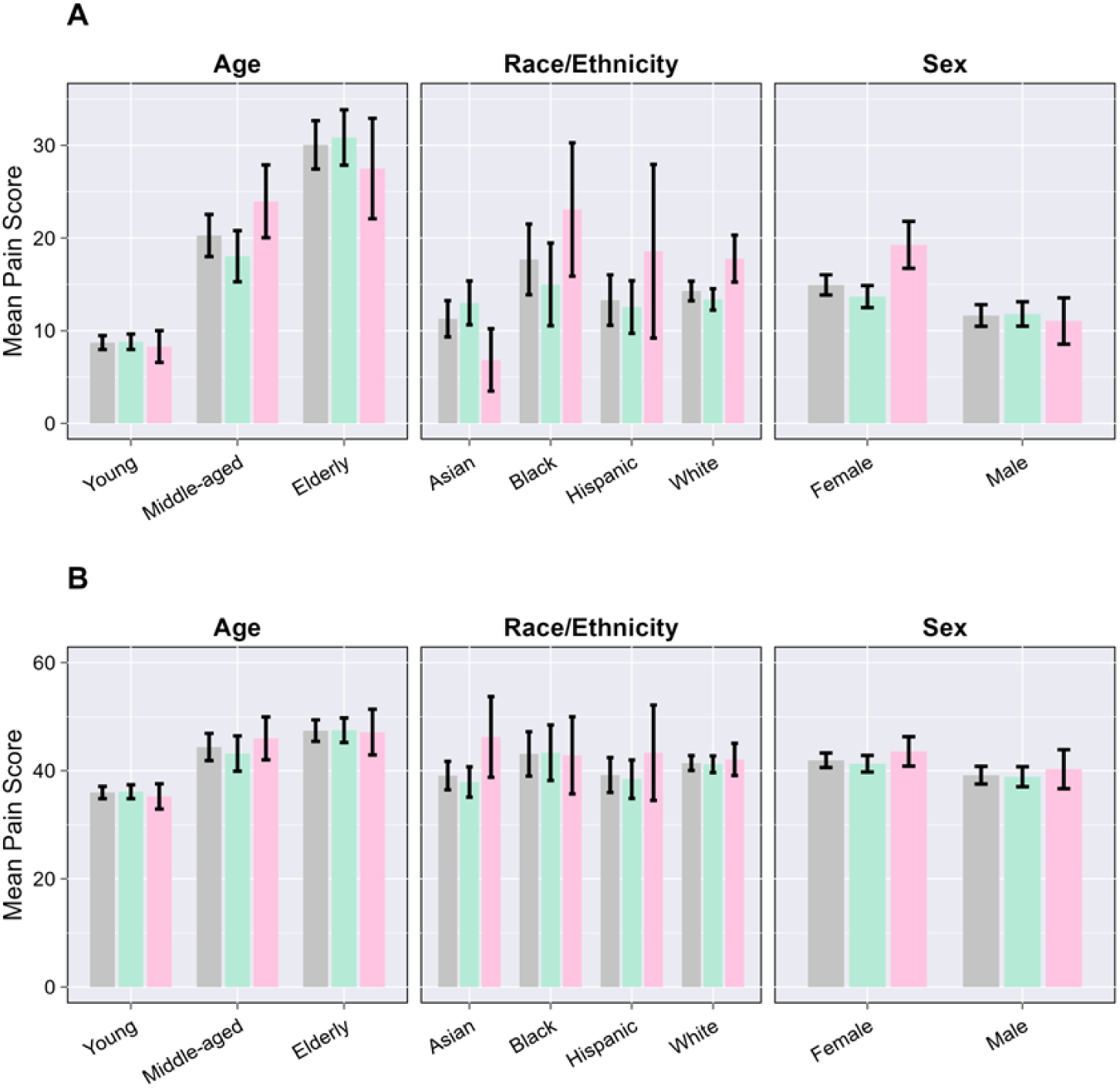
Average pain scores from (a) the primary analysis that included all participants and (b) the sensitivity analysis that included participants with pain scores greater than zero. Bars represent means across all participants (gray), participants with hand dominant hobbies (green), and participants with no hand dominant hobbies (pink) across the three demographic groups. Error bars indicate standard error.

The technical validation focused on contextualizing to what extent the BHaM population dataset represents known relationships between demographics, pain, and function. It importantly highlights limitations with generalizing population analyses to diverse populations and the need to carefully consider floor effects when performing analyses that include pain-related variables. Researchers are encouraged to keep these limitations in mind when using both the BHaM population dataset and the BHaM biomechanics dataset. Additionally, researchers should consider that the BHaM database includes biomechanics data from 29 different tasks and population data on hand strength, function, and dimensions. The inclusion of joint angles, muscle activations, external forces, and measured moments provide the necessary data for the best practices of validating musculoskeletal simulations.^22^ While MR images provide a physiologically-derived ground truth for developing new or personalized models. The inclusion of raw data within this published dataset allows fellow researchers to not be constrained by our processing methods. This means that inverse kinematics can be run with custom scripts outside OpenSim, analysis can be conducted on EMG frequencies, and other EMG normalization approaches can be implemented. Publishing this dataset is also one of the first steps towards developing an international standard for musculoskeletal modeling validations for the hand,^22^ and hopefully encourages the discussion of further contributions and standards for the musculoskeletal modeling community.

## 5 Code Availability

All R code scripts are publicly accessible on the project’s Kaggle repository.

## Acknowledgements

Thanks to the many members of the University of Florida’s Musculoskeletal Biomechanics Lab who assisted with data collection, including Sarah Barron, Chloe Baratta, and Tamara Ordonez Diaz. We also thank Judith Steadman for helping us obtain structural MR images from the upper limb. Thank you as well to Terrie Vasilopoulos for assistance with the statistical analyses described in the technical validation section of this manuscript. Funding from the National Institute of Biomedical Imaging and Bioengineering (Trailblazer Award R21 EB030068), the National Center for Advancing Translational Sciences (UL1TR001427), and National Science Foundation (Graduate Research Fellowship 1842473 and Cooperative Agreement DMR-2128556) is gratefully acknowledged.

## Author Contributions

J.B.H. and J.A.N. conceived the presented idea, secured funding, and oversaw all scientific and administrative aspects of the study. M.T.D. designed and conducted the experiments, processed the data and prepared it for publication, and wrote most of the manuscript. A.R.B. assisted with collection of the biomechanics dataset. T.F.K. conducted the statistical analyses and drafted the technical validation portion of the manuscript. K.M.K., E.M.L., I.T., and L.D. contributed to the experimental design of the study, including selection of the questions included in the population dataset and tasks included in the biomechanics dataset. W.S.B., J.B.N., and M.B.O. contributed to data processing, including analysis of hand images (W.S.B.) and preparing marker-based motion capture data for inverse kinematics (J.B.N. and M.B.O.). All authors assisted with collection of the population dataset and approved the final version of the manuscript.

